# Glucocorticoid and adrenergic receptor distribution across human organs and tissues: a map for stress transduction

**DOI:** 10.1101/2022.12.29.520757

**Authors:** Sophia Basarrate, Anna S. Monzel, Janell Smith, Anna Marsland, Caroline Trumpff, Martin Picard

**Author notes:** Address correspondence to Martin Picard, 1051 Riverside Dr. Kolb 4, Picard Lab New York, NY 10032.

## Abstract

**Objective:** Psychosocial stress is transduced into disease risk through energy-dependent release of hormones that affect target organs, tissues, and cells. The magnitude of the physiological stress responses reflects both systemic levels of these hormones and the sensitivity of target tissues to their effects. Thus, differential expression of receptors across organs likely contributes to stress transduction. Here we provide a quantitative whole-body map of glucocorticoid and adrenergic receptor expression.

**Methods:** We systematically examined gene expression levels for the glucocorticoid receptor (GR), α- and β-adrenergic receptors (AR-α1B, AR-α2B AR-β2, and AR-β3), across 55 different organs using the Human Protein Atlas dataset. We also leveraged the Human Proteome Map and MitoCarta3.0 data to examine receptor protein levels and, given the energy-dependence of the stress response, the link between stress hormone receptor density and mitochondrial pathways. Finally, we tested the functional interplay between GR activation and AR expression in living human cells.

**Results:** The GR was expressed ubiquitously across all investigated organ systems. Immune tissues and cells expressed the highest GR RNA and protein levels. In contrast, AR subtypes showed lower and more localized expression patterns. Co-regulation was found between GR and AR-α1B, as well as between AR-α1B and AR-α2B. In human fibroblasts, activating the GR selectively increased AR-β2 (3.6-fold) and AR-α1B (2.2-fold) expression, confirming their interaction. Consistent with the energetic cost of stress responses, GR and AR expression were positively associated with the expression of key mitochondrial pathways.

**Conclusion:** Our results provide a cartography of GR and AR expression across the human body. Tissue-specific stress hormone receptor expression patterns could make specific organ systems more responsive to the sustained, energetically expensive, neuroendocrine signaling pathways triggered by chronic psychosocial stress.

## Introduction

Stressful psychological experiences are transduced into physiological changes through the action of hormones on target cells and organs. The psychobiological stress response occurs particularly when an individual feels that environmental demands and exposures surpass their “adaptative capacity” (1). In the 1910s, Walter Cannon described the “emotional stimulation of adrenal secretion” – catecholamines – as a key endocrine factor in this brain-body axis (2). Subsequently, Hans Selye showed that stressful experiences triggered another kind of adrenal secretion, the peripheral release of the glucocorticoid hormones cortisol and corticosterone (3). Selye noted two hallmarks of stressful experiences: adrenal hyperactivity associated with increased release of glucocorticoids and catecholamines, and atrophy of lymphoid tissues of the immune system, which must have the ability to sense and respond to these stress signals. Over subsequent decades, the functional consequences of stress hormones on the immune system have been well described (4), including reliable changes in immune function and downstream consequences such as increased susceptibility to infections (5), slowed wound healing (6), and other psychoneuroimmunological processes that affect organs and tissues throughout the human body. All main organ systems are known to be acutely or chronically responsive to stress exposure, including the brain, heart, airways, liver, kidneys, and the primary/secondary immune tissues, possibly accounting for the association of life stress with increased risk for psychiatric disorders, cardiovascular disease, asthma, and immune-related conditions (7-10). However, not all organ systems are equally responsive to the activation of stress pathways. One factor that may underly differential responses of target organs to stress signals, and therefore the extent to which they are impacted by acute and chronic stress, is the variable expression of the receptors that sense and communicate stress signals across the organism.

The hypothalamic-pituitary-adrenal (HPA) and sympathetic-adrenal-medullary (SAM) axes operate with other stress systems to transduce psychosocial stress into peripheral physiological responses. Activation of these pathways results in the peripheral release of glucocorticoids and catecholamines, which bind to their respective receptors in or on target cells (7). The primary glucocorticoid hormone released in primates is cortisol, which binds to the glucocorticoid receptor (GR), encoded by the *NR3C1* gene. The GR is mainly located in the cytoplasm (although some is expressed on the plasma membrane) until activated by its glucocorticoid ligand, at which point it dimerizes and translocates into the nucleus and mitochondria where it acts as a transcription factor (11), modulating the expression of both nuclear and mitochondrial genes that catalyze a range of physiological responses (12, 13). The role of the GR in transducing psychosocial stress into biological changes has been documented extensively, particularly in the immune system and brain (14, 15).

The SAM axis acts in parallel with the HPA axis (7). The SAM axis rapidly in response to the experience of psychosocial stress by releasing the catecholamines epinephrine and norepinephrine from the adrenal medulla and sympathetic nerve terminals. Catecholamines bind to a broad family of G-protein coupled receptors located on cell membranes, termed adrenergic receptors (AR). The nine known human ARs are classified into α and β types, which are further divided into numbered subtypes (16, 17). The AR subtypes AR-α1B, AR-α2B, AR-β2, and AR-β3 (encoded by genes *ADRA1B, ADRA2B, ADRB2*, and *ADRB3*, respectively) have been shown to be involved in transducing psychosocial stress into biologically relevant effects (18-23).

Once adrenergic and glucocorticoid receptors transduce stress signals from the outside to the inside of target cells and tissues, this triggers energy-intensive molecular and physiological responses including increases in heart rate, vasodilation/constriction altering blood pressure, gene expression changes, and changes in neuronal excitability. Within cells, all of these hormone-mediated changes are powered by ATP generated through mitochondrial respiration (13, 24). Because mitochondria produce the energy and signals that enable the body to adapt to stress, there is a direct connection between stress responses and mitochondrial functions (24, 25). Mitochondria also contribute to the stress response by metabolizing catecholamines, in addition to generating the energy needed to power brain adaptations resulting from AR signaling during the stress response (26). On this basis, we can expect a functional connection between stress hormone signaling and mitochondrial regulation in target tissues, although this question has not been systematically examined in humans.

Thus, without receptors for glucocorticoids and catecholamines to convey messages carried by HPA and SAM axis signaling molecules, cells of the brain, immune system, digestive tract, and other organ systems would not be able to sense and respond to stressors. Variability in the density of these receptors across organs may help to identify systems that are likely to be more responsive and/or vulnerable to chronic stress. To explore the molecular basis for stress transduction across the human body, we leveraged publicly available datasets and live cell experiments to systematically quantify the RNA and protein expression of stress hormone receptors across human tissues, identify co-regulation between specific stress hormone receptor subtypes, and evaluate associations between GR and AR expression and mitochondrial gene expression. Our results provide a quantitative map of neuroendocrine stress hormone receptors across human tissues.

## Results

Our primary objective was to investigate differential expression of the GR and ARs that transduce psychosocial stress signals across human organs and their functional systems (Figure 1A). To accomplish this, we first analyzed transcriptomic (RNA sequencing) data from the *Human Protein Atlas* (27-29) and proteomic data from the *Human Proteome Map* (30), combining expression data across 55 different human tissues (n=948 individuals) and six immune cell subtypes (Supplementary Figure 1A). We confirmed the sensitivity of these data and our ability to discriminate between ubiquitously expressed genes, such as the cytoskeletal element actin (*ACTB*) present in every nucleated cell of the body, and genes with specialized expression restricted to some tissues such as the serotonin receptor (*HT2RA*) in neurons among the central nervous system (CNS), or the B cell marker CD19 expressed exclusively in immune tissues and to some extent in organs known to harbor tissue-resident immune cells (Supplementary Figure 1B). The results confirmed our ability to sensitively discriminate genes expressed either ubiquitously throughout the human body or restricted to only one or a few tissues.

**Figure 1.**
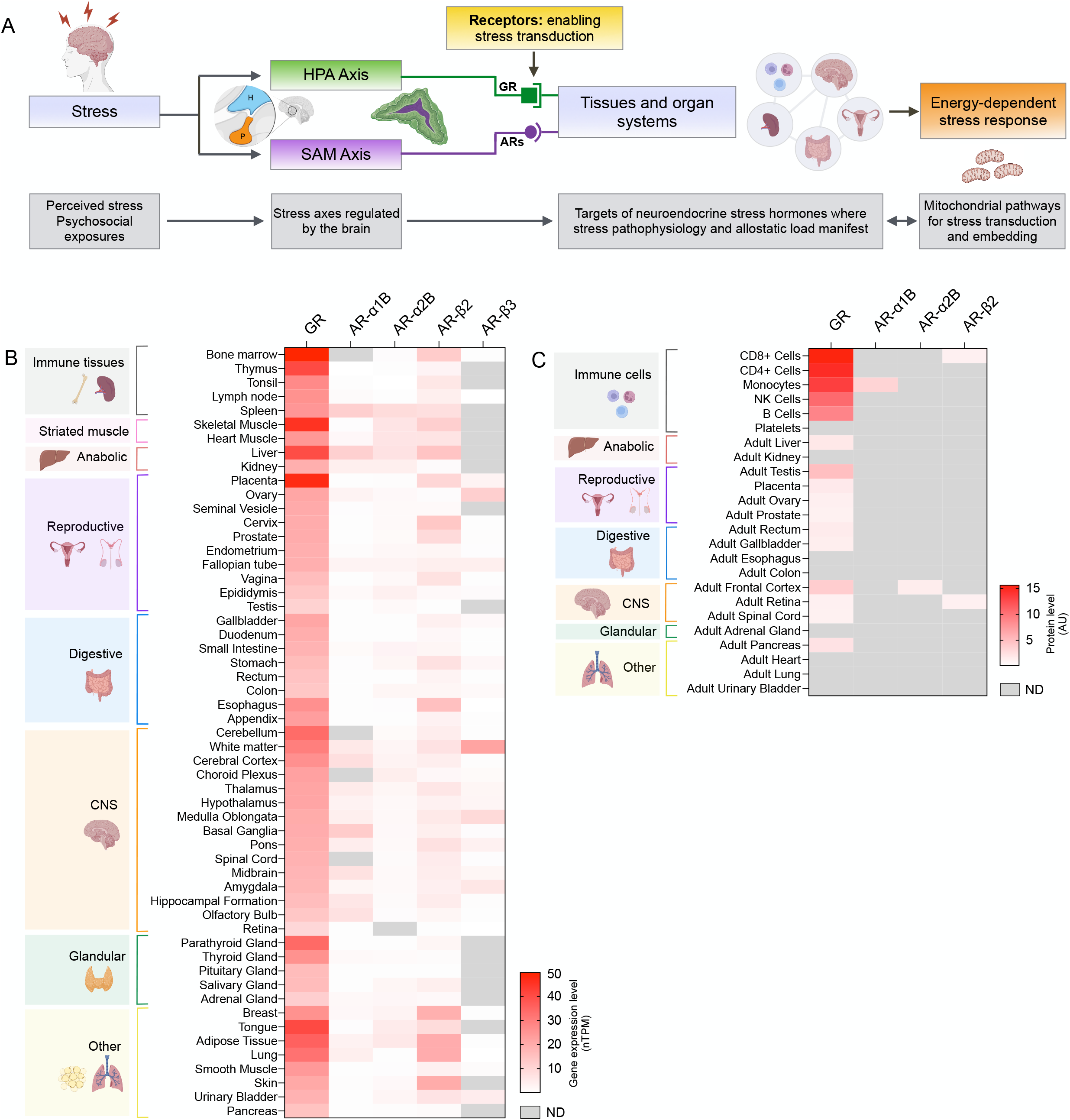
Stress hormone receptors transduce experiences of psychosocial stress to tissues; heatmaps of glucocorticoid and adrenergic receptor RNA and protein expression. (**A**) The two major pathways with receptors enabling the transduction of psychosocial stress are illustrated here: the HPA and the SAM axes. The signaling hormones from these axes act on target organs via glucocorticoid (GR) and adrenergic receptors (ARs), which mobilizes energy-dependent responses sustained by mitochondrial energy production and signaling. (**B**) Heatmap of glucocorticoid and adrenergic receptor RNA expression (in normalized transcript per million [nTPM]) in different human tissue types (n=55), grouped by organ system. (**C**) Heatmap of glucocorticoid and adrenergic receptor protein expression (in arbitrary units [AU]) across tissue types (n= 17 adult tissues, n=8 immune cell types), grouped by organ system (Uhlén M et al., 2015; Kim et al., 2015). *Abbreviations*: *ARs*, adrenergic receptors; *AR-α1B*, adrenergic receptor alpha 1B; *AR-α2B*, adrenergic receptor alpha 2B; *AR-β2*, adrenergic receptor beta 2; *AR-β3*, adrenergic receptor beta 3; *CNS*, central nervous system; *GR*, glucocorticoid receptor; *HPA*, hypothalamic pituitary adrenal; *ND*, not detectable; *SAM*, sympatho-adreno-medullary.

### GR and ARs are expressed heterogeneously across organ systems

Our systematic survey showed that GR is expressed ubiquitously across all investigated organ systems, theoretically enabling every human tissue to respond to cortisol (Figure 1B). Comparing organ systems to one another, striated muscles had the highest GR expression (average expression of 34.9 normalized RNA transcript per million [nTPM]) consistent with the role of glucocorticoids in muscle metabolism (31). The immune system exhibited the second highest average GR expression, 64.4% higher than the central nervous system (CNS) (Hedge’s g=1.70, p<0.01, independent sample T-test, Welch correction), digestive (g=1.83, p<0.05), and reproductive organs (g=1.18, p<0.05). Among individual tissues, the bone marrow (49.8 nTPM) and the placenta (47.3 nTPM) contained the highest GR expression, consistent with the well-described sensitivity of the immune system and placenta to glucocorticoid signaling (32).

Relative to GR, AR subtypes expression across all organ systems showed on average 9.8-fold lower expression (p<0.0001, ANOVA). In relation to α (alpha) ARs subtypes, AR-α1B gene expression was on average highest in anabolic tissues, with especially high levels detected in the liver, where catecholamines are well known to stimulate gluconeogenesis (33). The CNS also had high AR-α1B RNA levels, particularly in the basal ganglia and cerebral cortex. AR-α2B transcripts were detected at low levels in all tissue types, except the retina, and its expression was highest in the spleen, liver, and striated muscles, relatively consistent with GR.

In relation to β (beta) ARs subtypes, AR-β2 transcripts were expressed ubiquitously, although at lower levels than GR (3.9-fold lower on average). In contrast to α ARs, AR-β2 had the highest expression in the skin, lung, and adipose tissue. AR-β3 was only expressed in 69% of tissues, although it was expressed at high levels in some isolated brain regions, including white matter tracts and the medulla, as well as the ovaries.

The gene expression levels of all stress hormone receptors in each tissue type are detailed in Supplementary Table 1, along with the additive expression of the five examined GR and ARs. These results emphasize the particularly robust expression of stress hormone receptors in metabolic, CNS, and immune tissues, consistent with the known energetic recalibrations that these organ systems undergo in response to acute stress (34).

### GR and AR receptor protein abundance

To be functionally active, the RNA transcripts discussed above must be translated into protein receptors at the cell surface. Thus, although protein abundance is technically more difficult to measure than RNA, it represents the most direct assessment of a tissue’s molecular sensing machinery. Similar to patterns observed for RNA, the GR protein was detected in the largest variety of tissues, including all reproductive and CNS tissues (Figure 1C). GR was detected at highest levels in immune cells (except platelets), providing the molecular basis for glucocorticoid sensitivity of the immune system. Compared to GR, all ARs showed significantly lower and more restricted protein expression, consistent with patterns observed at the RNA level. Compared to average GR protein levels, AR-α1B was 96% lower, AR-α2B was 98% lower, and AR-β2 was 97% lower. AR-α1B protein was only detected in monocytes, while AR-α2B was only detected in the adult frontal cortex. AR-β2 protein was expressed in CD8^+^ T cells and in the adult retina. AR-β3, likely of very low abundance, was not detected in any of the examined tissues. The protein abundance for all examined receptors and tissues are detailed in Supplementary Table 2.

### GR and ARs co-regulation across tissues

Under some conditions, psychosocial stress differentially activates both the HPA and SAM axes, with SAM responses occurring more rapidly than HPA responses. We next examined if tissues that express more GRs also express more ARs. Our correlation-based co-regulation analysis of gene expression across 55 tissues showed that organs that expressed higher levels of GR transcripts also generally expressed higher levels of AR-β2 receptor RNA (Spearman r=0.50, p=0.0009, Bonferroni-corrected), but not other ARs (Figure 2A). Tissues like the bone marrow, skin, and placenta express high levels of both receptors, potentially providing particularly high sensitivity of these tissues to glucocorticoids and catecholamines (Figure 2B). Although less robust, AR-α1B and AR-α2B also showed a significant correlation (r=0.46, p=0.004, Bonferroni-corrected), with the spleen and liver expressing particularly high levels of both receptors (Figure 2C).

**Figure 2.**
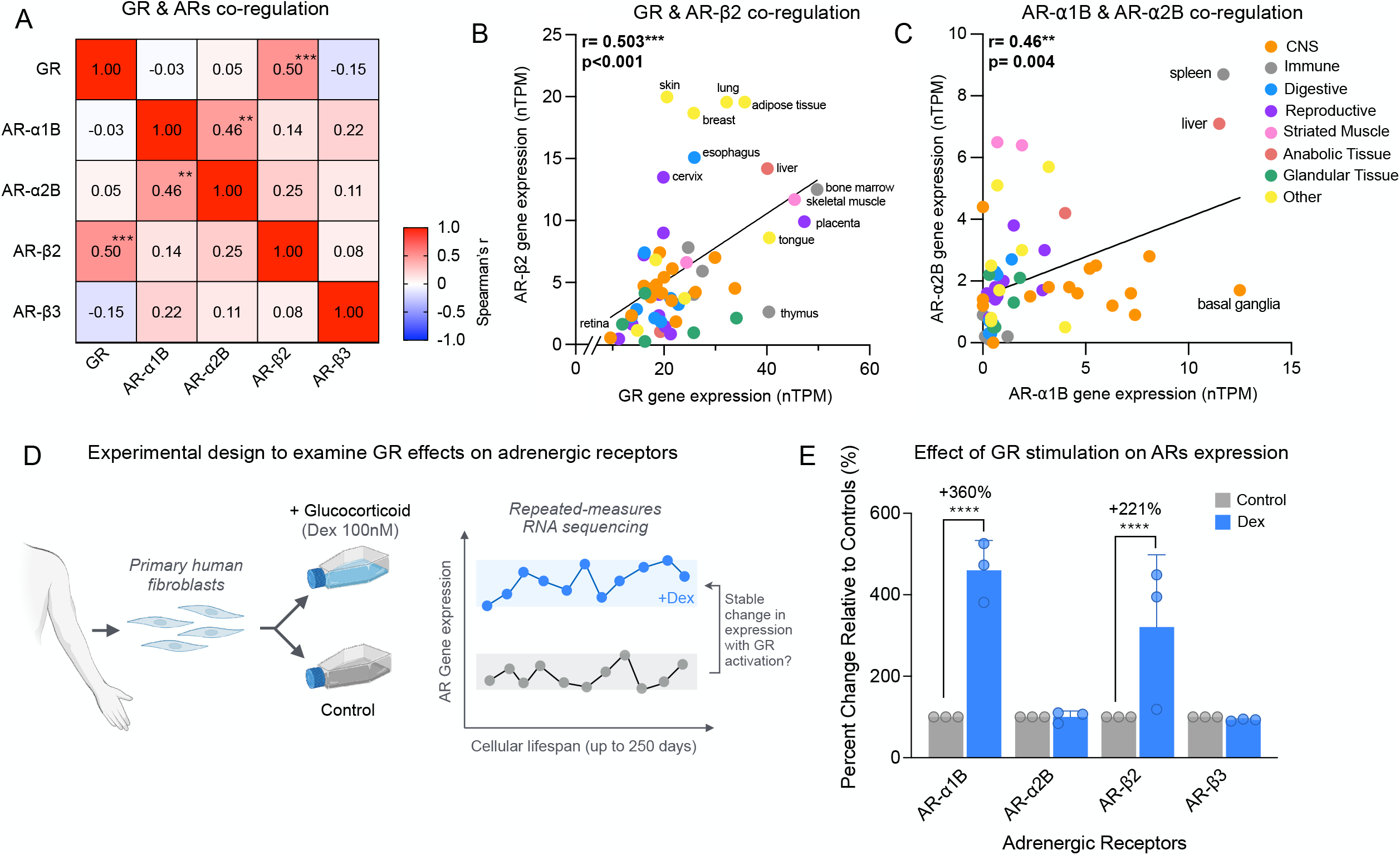
Co-regulation and functional interaction of stress hormone receptor expression. (**A**) Correlation matrix of glucocorticoid and adrenergic receptor RNA expression. (**B**) Scatterplot of GR and AR-β2 RNA expression, which showed a significant positive correlation. (**C**) Scatterplot of AR-α1B and AR-α2B, which showed a significant positive correlation. (**D**) Schematic of study design where primary dermal fibroblasts are obtained from forearm and cultured *in vitro*, followed by longitudinal gene expression analysis across the cellular lifespan. The average of all timepoints for each donor was taken as a measure of stable lifetime gene expression for each adrenergic receptor (AR), compared to the values in the untreated (control) culture of the same donor. (**E**) Effect of Dex (Dex, 100nM) on the gene expression of ARs in n=3 fibroblast lines (healthy control 1, 2, 3). Data are means ± SEM, normalized to control. Statistics from Spearman’s rank correlation with Bonferroni correction (A, B), and mixed effects model (D), p<0.05*, p<0.01**, p<0.001***, p<0.0001****. *Abbreviations: AR*, adrenergic receptor; *AR-α1B*, adrenergic receptor alpha 1B; *AR-α2B*, adrenergic receptor alpha 2B; *AR-β2*, adrenergic receptor beta 2; *AR-β3*, adrenergic receptor beta 3; *Dex*, dexamethasone; *GR*, glucocorticoid receptor.

### GR signaling upregulates adrenergic receptor expression in vitro

To examine if the co-regulation of GR and ARs may reflect functional coupling at the transcriptional level, we leveraged an *in vitro* gene expression (RNA sequencing) dataset of primary human fibroblasts treated with the GR agonist dexamethasone (Dex), a synthetic glucocorticoid (Figure 2D) (35). In this cellular *in vitro* system, as in most human tissues, GR was the most highly expressed receptor, and all ARs were expressed at considerably lower levels. This result and other molecular features, such as DNA methylation patterns (36) mirror measurements in human tissues, providing evidence of external validity for this *in vitro* cellular system.

In line with our findings in the human body, experimentally inducing GR signaling increased expression levels for two of the four ARs examined and has no effect on two (Figure 2E). Dex induced AR-α1B expression by 360% (p<0.0001, mixed effects model) and AR-β2 expression by 221% (p<0.0001). The functional induction of AR-β2 by GR activation corroborates and supports the *in vivo* evidence that GR and AR-β2 are co-regulated across human tissues. This finding also is consistent with a functional interplay among the HPA and SAM axes at the level of target tissues.

### AR-α1B and AR-α2B transcripts are positively associated with mitochondrial gene expression

Given that the stress response consumes energy mostly provided by mitochondria (25), we systematically assessed the relationship between GR and AR receptors and all known mitochondrial genes in the tissue gene expression data (27-29). Mitochondrial content was estimated for each tissue type by taking the average gene expression of all mitochondrial genes, based on the MitoCarta3.0 database (34, 37). Estimated tissue mitochondrial content was positively associated with AR-α1B (Spearman r=0.47, p= 0.0003, Bonferroni corrected) and with AR-α2B gene expression (r=0.55, p=0.0009, Bonferroni corrected) (Figures 3A and 3C), indicating that tissues with more of these adrenergic receptors express mitochondrial genes at higher levels.

**Figure 3.**
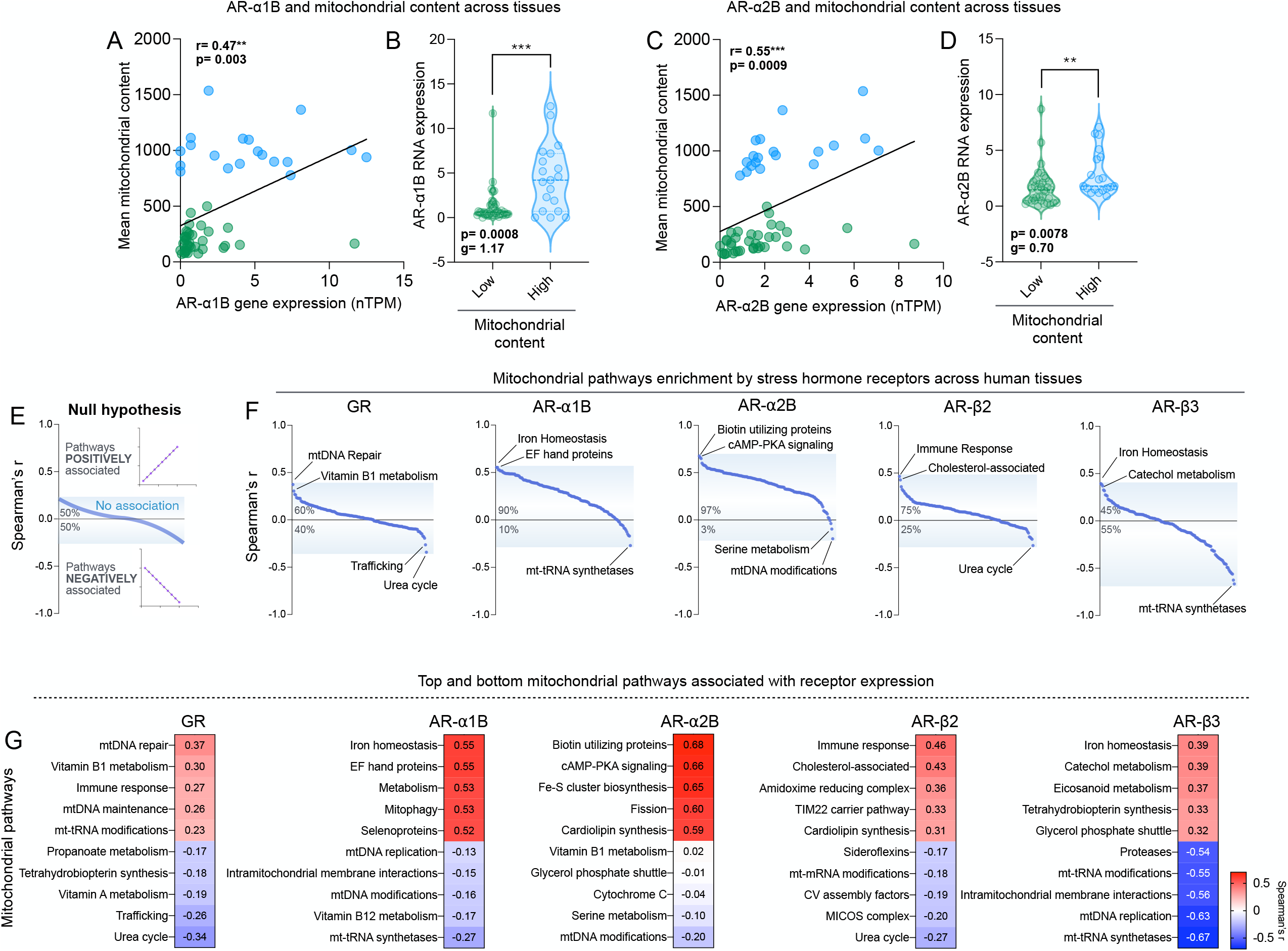
Stress hormone receptor and mitochondrial protein RNA expression. (**A**) Scatterplot significant correlation (Spearman’s r) between AR-α1B receptor RNA expression (nTPM) and mean mitochondrial content (mean nTPM). (**B**) Bar graph of average AR-α1B RNA expression for tissues dichotomized by high versus low mean mitochondrial content (threshold= 500 nTPM). (**C, D**) Same as in A and B, for AR-α2B receptor. (**E**) Null hypothesis plot of correlations (Spearman’s r) between stress hormone gene expression and RNA expression of mitochondrial proteins involved in specific functional pathways (**F**) Plot of correlations (Spearman’s r) between stress hormone receptor gene expression and RNA expression of mitochondrial pathway proteins. (**G**) Heatmaps of five most positive and most negative correlations between stress hormone receptors and mitochondrial pathway gene expression. Statistics from Mann-Whitney test, p<0.05*, p<0.01**, p<0.001***, p<0.0001****. *Abbreviations: AR-α1B*, adrenergic receptor alpha 1B; *AR-α2B*, adrenergic receptor alpha 2B; *AR-β2*, adrenergic receptor beta 2; *AR-β3*, adrenergic receptor beta 3; *nTPM*, normalized transcripts per million; *GR*, glucocorticoid receptor.

The distributions of our index of mitochondrial content for AR-α1B and AR-α2B were bimodal, indicating two main groups of tissues with either low or high mitochondrial gene expression. We therefore took a different approach, comparing AR-α1B and AR-α2B expression levels between the two groups of tissues (Figures 3B and 3D). Consistent with the correlation analysis, tissues with higher estimated mitochondrial content, such as the heart muscle, cerebral cortex, skeletal muscle, and liver, had higher expression of both AR-α1B and AR-α2B. Thus, tissues endowed with high stress hormone receptor density may harbor commensurately higher capacity for energy production and other mitochondrial functions.

### Stress hormone receptors are associated with specific mitochondrial pathways

Mitochondria perform multiple functions beyond energy transformation including vitamin metabolism, nucleotide synthesis for DNA replication and telomere maintenance, hormone and neurotransmitter synthesis, reactive oxygen species (ROS) production, and others (38). From the point of view of the cell, mitochondria sit at the interface of the external environment and the internal (epi)genome. Many of these mitochondrial functions are likely responsible for transducing stressors and the resulting neuroendocrine signals into the molecular changes that contribute to stress pathophysiology (39). Therefore, defining the mitochondrial pathways most strongly (or weakly) associated with GR and AR expression may provide insights or future targets to understand the psychobiological mechanisms responsible for the biological embedding of psychosocial stress.

To systematically examine whether the expression of GR and AR receptors was associated with specific mitochondrial pathways, we correlated the five stress hormone receptors observed across 55 human tissues with each of the 149 well-defined functional pathways in MitoCarta3.0 (34). For each receptor, this yielded 149 regression coefficients, which we ranked from the highest to lowest. For this analysis, if there is no association between the expression of GR/ARs and mitochondrial pathways, the distribution would be centered around zero and follow a Gaussian distribution (i.e., small positive and negative tails, reflecting the null hypothesis) (Figure 3E). The results showed that for all receptors except AR-β3, the Spearman’s Rho distributions were positively skewed. There were significantly more mitochondrial pathways positively associated with GR and ARs than expected by chance (45-97%, average 81% vs 50% expected by chance, p<0.0001, Chi square) (Figure 3F), suggesting that tissues with high sensitivity to glucocorticoids and catecholamines express high levels of genes associated with various mitochondrial pathways.

The identified mitochondrial pathways included some expected, as well as some novel stress-transduction mitochondrial profiles. Figure 3F illustrates the distributions, and Figure 3G lists the top and bottom 5 mitochondrial pathways associated with each receptor. Overall, there was little overlap between the mitochondrial profiles associated with each receptor (Supplemental Figure S2), suggesting either the existence of highly receptor-specific mitochondrial profiles, or the influence of confounding factors and low signal-to-noise ratio for these analyses. Here we highlight some relevant patterns.

Consistent with the role of calcium signaling downstream from adrenergic receptors, tissues with high AR-α1B were enriched for mitochondrial *EF hand proteins*, and AR-α2B was enriched for mitochondrial *cyclic AMP-protein kinase A (cAMP-PKA) signaling*, which represent canonical intracellular calcium-related signaling systems. Specifically related to mitochondria, high AR-α2B and AR-β2 were linked to *cardiolipin synthesis*, a specialized mitochondrial phospholipid essential for normal function. High GR was linked to mitochondrial DNA (*mtDNA) repair*, which safeguards mtDNA genes required to sustain energy transformation, in addition to *mtDNA maintenance*. This result is consistent with the energetic cost of stress (40), and perhaps also with an emerging literature around the role of mtDNA signaling in response to acute and chronic stress (41). AR-β3 and GR were also positively associated with the mitochondrial *immune response* pathway, which could reflect the enhancement of some parts of the immune system in response to stress (42). Overall, these data reveal predominantly positive associations between the expression of stress hormone receptors and mitochondrial pathways, consistent with the notion that HPA and SMA stress signaling entails energy costs at the cellular level.

## Discussion

To map the signaling machinery responsible for the transduction of psychosocial stress into coordinated physiological responses, we evaluated the RNA and protein expression of glucocorticoid and adrenergic receptors across a wide range of organ systems. Our results demonstrate a few important points. First, in contrast to ARs that are expressed at lower levels and in a more tissue-specific pattern, the GR is ubiquitously expressed and at considerably higher levels across organ systems. Second, body-wide and live-cell analyses show that GR and some ARs are co-regulated, consistent with their functional and genetic interactions. Finally, in line with the energetic cost of stress responses, we find evidence that tissues with greater GR and ARs density, and presumably greater sensitivity, harbor higher estimated mitochondrial content and expression of specific mitochondrial pathways. These findings provide a quantitative body-wide inventory of receptors involved in brain-body communication and may help to understand how acute and chronic stress is transduced differentially across human organ systems.

The distribution of GR and ARs across the body is the mechanism by which psychobiological stress is transduced into cellular responses to activation of the HPA and SAM axes. As expected, organ systems classically implicated in response to HPA activation have the highest GR levels, making them particularly sensitive and responsive to circulating glucocorticoids (43). These include the *liver*, which breaks down glycogen stores and performs gluconeogenesis to increase blood glucose, the *skeletal muscles*, where glucocorticoids suppress glucose uptake to increase availability to other organs like the *brain*, and the adaptive *immune system*, which similarly must be rapidly suppressed to prevent overconsumption of glucose and other circulating energy substrates in times of stress. All immune organs and purified immune cells showed high levels of GR, especially at the protein level, which may enable particularly rapid and robust responses to HPA signaling. On the other hand, high receptor abundance also may render them more vulnerable to adaptation arising from HPA axis overactivation during chronic stress. The hypersensitivity of immune organs to stress signaling aligns with the large body of research documenting stress effects on various aspects of immunity, as evidenced by down-regulation of aspects of adaptive immune function and increased susceptibility to the common cold (5, 44-46). Additionally, we found that the placenta exhibits substantial GR expression, indicating a high potential for GR signaling to mediate the prenatal effects and vulnerability of fetal development to maternal stress exposure (47).

Compared to GR, the lower and more localized protein expression of AR subtypes suggests that the organs responsive to stress signals conveyed by the SAM axis are more specialized or selective. For example, the AR-α1B protein was only detected in monocytes, which aligns with previous findings about the influence of AR-α1 on innate immune cells (48, 49). Additionally, AR-α2B protein was only identified in the adult frontal cortex, potentially rendering this brain region particularly sensitive to local or systemic catecholaminergic signals arising from psychosocial stress signals. These selected examples of more circumscribed AR expression may provide a biological explanation for organ-specific responses to adrenergic stimulation. However, given that the fight or flight response is known to involve the swift, coordinated reaction of a range of organs, it is also possible that the AR protein levels were simply below the detection threshold in some tissues.

Stress signaling axes do not act in isolation. Our findings that GR and AR-β2 are co-regulated across human tissues could suggest that some organs are sensitive to both HPA and SAM axis outputs. The high levels of both receptors particularly in bone marrow, placenta, liver, adipose tissue, and skin could enable either more robust or faster responses in these tissues, as well as their coordination. Moreover, our *in vitro* results demonstrating that GR stimulation increases the expression of ARs (α1B and β2) confirms a potential functional connection between these pathways, where the presence of glucocorticoids leads to sensitization to catecholamine signaling, at least in some tissues.

Finally, our findings established an initial connection between GR, ARs, and mitochondrial gene expression among human tissues. The significant correlation of levels of both AR-α1B and AR-α2B with estimated mitochondrial content indicate that the tissues most sensitive to adrenergic stimulation may be better prepared to mount robust energetic responses. Additionally, the expression of mitochondrial genes involved in the cAMP-PKA signaling pathway showed a strong positive association with AR-α2B. Given that ARs are G-protein coupled receptors that typically rely on cAMP-signaling pathways, the high expression of these mitochondrial pathways could enable faster or more efficient intracellular communication in these tissues, thereby contributing to more energetically efficient stress responses (16). Additionally, the positive correlation of GR transcript expression with mtDNA repair-related mitochondrial genes suggests that stress-susceptible tissues are also preparing to repair mtDNA damage that may result from stress (50). However, there were negative associations between AR-β3 and various mitochondrial pathways, indicating that tissues sensitive to stress transduced by this receptor may predominantly activate other intracellular pathways less dependent upon mitochondria. Overall, the diverse associations between stress hormone receptor expression and a wide range of mitochondrial pathways empirically supports the general conceptual link between stress responses and mitochondrial biology (51). These preliminary results identify potential candidate pathways to examine the cellular mechanisms responsible for the biological embedding of trauma, adversity, and chronic stress (52), which call for further targeted studies.

One limitation of this work leveraging gene expression is the indirect assessment of the effect of stress on tissues, as this method does not capture the dynamic transcription, translation, and degradation of stress hormone receptors over time. Additionally, the *Human Protein Atlas* and *Human Proteome Map* datasets did not include RNA and protein expression in the exact same set of organs and immune cells, allowing for less direct comparison between the two (RNA and protein) modalities. Also, the *Human Proteome Map* did not include protein levels for AR-β3, likely undetected because of its low expression level. Technically, the untargeted proteomics data was generated by mass spectrometry, which has a fairly high detection limit and may not have detected the proteins present at low levels (53). A more sensitive method, such as fluorescent peptide fingerprinting or other targeted approaches with finer spatial resolution could be more suitable to precisely quantify specific proteins in organs of interest (53). We also note that our whole-body transcriptomic data reflect aggregate data across ∼950 participants. Although this adds confidence that the results are generally robust, it does not permit the examination of inter-individual differences. Whether individuals exhibit differences in GR and ARs distribution that could contribute to the magnitude of the physiological stress response should be considered in future studies. Additionally, single-cell studies of GR and AR expression are needed given the differential effects of stress on distinct cell types, such as those of the innate versus adaptive immune system, which likely co-exist within the immune tissues and organs sampled. Finally, some tissues showed no stress hormone receptor expression, such as platelets, despite extensive literature highlighting the way in which they are impacted by stress (54). This calls for further consideration of downstream effects of the HPA and SAM axes, as well as non-classical mechanisms of stress transduction beyond these two well-studied types of receptors.

In conclusion, this study provides a rough cartography of canonical glucocorticoid and adrenergic receptors, highlighting their high heterogeneity among organs across the human body. The resulting quantitative map of organ-specific stress hormone receptor expression (available in Supplemental Tables 1-2) provides a basis to understand the nature and magnitude of stress responses among specific organ systems, such as the immune system, that occur following the activation of neuroendocrine pathways by psychosocial stress. This analysis also demonstrates a connection between GR and adrenergic receptors, as well as with mitochondrial gene expression, consistent with the notion that mitochondria and cellular energetics contribute to transduce stressful experiences into physiological responses relevant to human health.

## Methods

### Data extraction

Stress hormone receptors were selected based on evidence linking specific receptor types to the transduction of psychosocial stress (18-22). Using the Human Protein Atlas, we systematically examined gene expression levels (RNA) for the glucocorticoid receptor (GR), α-adrenergic receptors (AR-α1B and AR-α2B), and β-adrenergic receptors (AR-β2 and AR-β3) across different organs (n=55) (55). For RNA expression, we utilized the Human Protein Atlas consensus tissue gene dataset with consensus normalized transcriptomics data determined from the HPA RNA-seq dataset (n= 375 normal tissue samples) and the Genotype-Tissue Expression (GTEx) Project RNA-Seq dataset (n= 948 donors, n= 17,382 total samples) (27, 56, 57). Human Protein Atlas data can be downloaded at https://www.proteinatlas.org/about/download.

For our analyses, organs were grouped into systems based on functional similarities, including the immune (n=5, e.g., spleen, lymph nodes, thymus), brain and central nervous system (CNS, n= 15, e.g., amygdala, cerebellum, cortex), digestive (n= 7, e.g., e.g., stomach, colon, duodenum), reproductive (n= 11, e.g., ovary, cervix, testis), and other systems (n= 17). For stress hormone receptor protein expression, we utilized the Human Proteome Map dataset with protein expression levels determined by high-resolution Fourier-transform mass spectrometry for adult tissues (n=18) and immune cell subtypes (n= 6) (30). Samples were pooled from three individuals before analysis(30). AR-β3 protein data was not detected. Human Proteome Map data can be downloaded at http://www.humanproteomemap.org/download.php.

### Cellular lifespan gene expression analysis to glucocorticoid stimulation

The glucocorticoid-treated fibroblast gene expression data was analyzed from (35, 50). Briefly, primary dermal fibroblasts harvested from 3 unrelated healthy donors were cultured throughout their replicative lifespan, passaging every ∼5 days either in untreated condition or chronically treated with the GR agonist dexamethasone (100nM). Cells were collected and RNA was isolated at 8-11 timepoints for each donor and used for RNA sequencing. The data was processed and expressed as normalized transcript per million (nTPM) as described in (35). To obtain a stable estimate of ARs gene expression for each donor, target ARs nTPM were averaged across all timepoints.

### Analyses of mitochondrial content and functional pathways

To extract mitochondrial genes, the *Human Protein Atlas* normalized tissue consensus dataset (27, 29) was mapped to the human MitoCarta3.0 (34) gene list (1136 mitochondrial genes). For each tissue, a transcription-based index of mitochondrial content was calculated from the average expression of all normalized (normalized transcripts per million, nTPM) mitochondrial transcripts. In addition, a score for MitoCarta-annotated mitochondrial pathways (149 pathways) was calculated for each tissue and used in regression analyses with each stress hormone receptor. First, the specific gene set of each pathway was extracted. Next, a tissue-specific unweighted gene expression score was calculated by taking the average expression of the specific mitochondrial gene set of each pathway.

### Statistical analyses

Organ systems-level RNA differences were quantified using Hedge’s g, a standardized measure of effect size, which guards against bias from small sample sizes (58). Independent samples t-tests were used to compare average gene expression of GR in different organ systems (CNS vs immune, CNS vs digestive, CNS vs reproductive). The Welch’s correction was applied for the comparison of GR expression in CNS and digestive tissues because the standard deviations differed significantly. Because of their exploratory nature, the t-test p-values were not adjusted for multiple comparisons. Non-parametric correlation (Spearman’s r) was performed to assess co-regulation between stress hormone receptors’ RNA expression and between mitochondrial pathway scores and stress hormone receptor RNA expression, using the Bonferroni correction to adjust for multiple comparisons. The cutoff for the high and low mitochondrial content groups was set at 500 nTPM, and a non-parametric Mann-Whitney U test was conducted to evaluate between-group differences. The effect of GR stimulation on ARs gene expression in human fibroblasts was tested using a mixed-effects model, using all timepoints available across the lifespan of each cell line to derive a stable estimate of expression level for each gene. Chi-square tests were used to compare the observed proportions of positive and negative correlations between mitochondrial pathways and stress hormone receptor expression compared to the proportions expected by chance (50:50). All analyses were performed in Prism (version 9), Excel (version 16.59), and R version 4.2.0 (2022-04-22) -- “Vigorous Calisthenics”.

## Abbreviations

*AR*: adrenergic receptor
*AR-α1B*: adrenergic receptor alpha 1B
*AR-α2B*: adrenergic receptor alpha 2B
*AR-β2*: adrenergic receptor beta 2
*AR-β3*: adrenergic receptor beta 3
*CNS*: central nervous system
*Dex*: dexamethasone
*GR*: glucocorticoid receptor
*HPA*: hypothalamic-pituitary-adrenal
*nTPM*: normalized transcripts per million
*SAM*: sympathetic-adrenal-medullary
*mtDNA*: mitochondrial DNA

## Acknowledgements

The authors are grateful to Gabriel Sturm for advising on analyses related to the Cellular Lifespan Dataset, and to the investigative GTEx and HPA teams that generated the whole-body RNAseq and proteomic datasets that made possible this project.

## SUPPLEMENTAL MATERIAL

**Figure S1.**
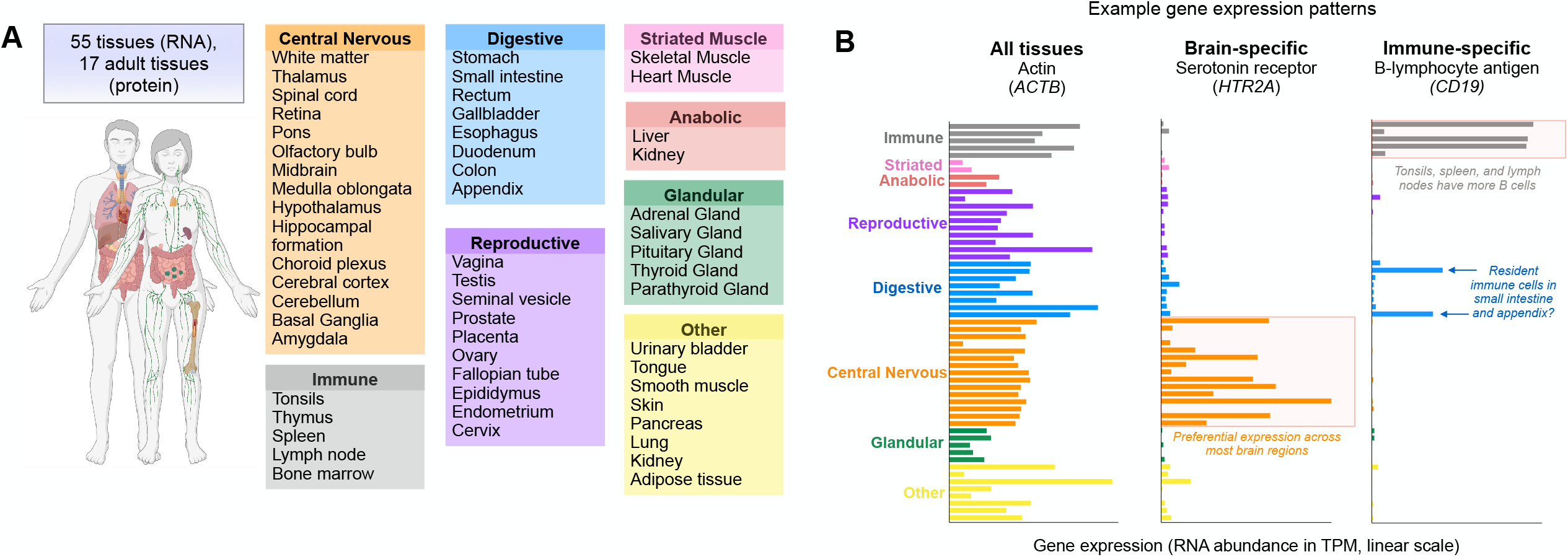
Overview of study design. (**A**) Study Design: Human Protein Atlas RNA immunohistochemistry data was acquired and categorized into major organ systems: central nervous system (n=15), immune system (n=5), reproductive system (n=10), digestive system (n=8), striated muscle (n=2), anabolic tissues (n=2), glandular tissues (n=5), and other organs (n=8) (Uhlén M et al., 2015). Mass spectrometry protein data for adult tissues (n=17) and immune cells (n=8) was acquired from Human Proteome Map (Kim et al., 2015). (**B**) Examples of genes ubiquitously expressed across organ systems (*ACTB*, actin beta protein), show specialized expression in the brain (*HTR2A*, serotonin receptor), or expressed specifically in immune system tissues (*CD19*, B-lymphocyte antigen).

**Figure S2.**
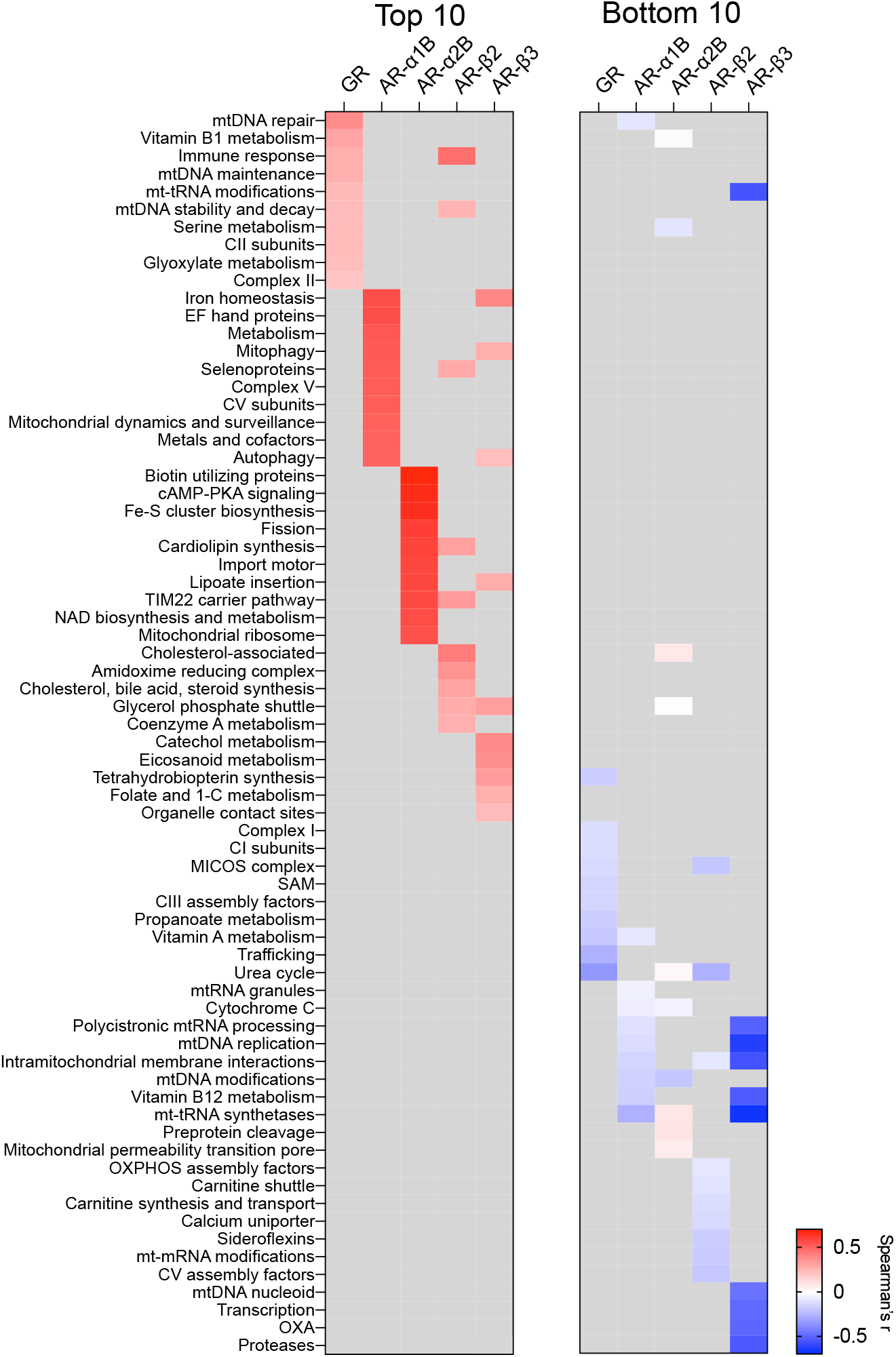
Correlations between stress hormone receptor expression and mitochondrial function pathway gene expression. Heatmap of (Spearman’s rank) correlations between RNA expression stress hormone receptor proteins and proteins associated with mitochondrial pathways. The mitochondrial pathways with the 10 highest (*left*) and 10 lowest (*right*) correlations are shown.

**Table S1 and Table S2. Stress Hormone Receptors Supplemental Tables.xlsx**

## Notes

**Conflicts of Interest and Source of Funding:** Work of the authors was supported by NIH grants R01MH119336, R01MH122706, R01AG066828, the Wharton Fund, and the Baszucki Brain Research Fund to M.P. The authors report no conflicts of interest.

http://www.humanproteomemap.org/download.php

https://www.proteinatlas.org/about/download

https://www.broadinstitute.org/mitocarta/mitocarta30-inventory-mammalian-mitochondrial-proteins-and-pathways

### Competing Interest Statement

The authors have declared no competing interest.

## References

1. S. Cohen, R. C. Kessler, L. U. Gordon, Measuring Stress: A Guide for Health and Social Scientists (Oxford University Press, 1997).

2. W. B. Cannon, D. d. l. Paz, Emotional Stimulation of Adrenal Secretion. American Journal of Physiology-Legacy Content 28, 64–70 (1911).

3. H. Selye, A Syndrome produced by Diverse Nocuous Agents. Nature 138, 32–32 (1936).

4. J. I. Webster Marketon, R. Glaser Stress hormones and immune function. Cell Immunol 252, 16–26 (2008).

5. S. Cohen, D. A. Tyrrell, A. P. Smith, Psychological stress and susceptibility to the common cold. N Engl J Med 325, 606–612 (1991).

6. J. P. Gouin, J. K. Kiecolt-Glaser, The impact of psychological stress on wound healing: methods and mechanisms. Immunol Allergy Clin North Am 31, 81–93 (2011).

7. G. Russell, S. Lightman, The human stress response. Nature Reviews Endocrinology 15, 525–534 (2019).

8. C. Tsigos, I. Kyrou, E. Kassi, G. P. Chrousos, “Stress: Endocrine Physiology and Pathophysiology” in Endotext, K. R. Feingold et al., Eds. (MDText.com, Inc., South Dartmouth (MA), 2000).

9. R. H. Straub, M. Cutolo, Psychoneuroimmunology-developments in stress research. Wien Med Wochenschr 168, 76–84 (2018).

10. M. R. Irwin, S. W. Cole, Reciprocal regulation of the neural and innate immune systems. Nature Reviews Immunology 11, 625–632 (2011).

11. A.-M. G. Psarra, C. E. Sekeris, Glucocorticoids induce mitochondrial gene transcription in HepG2 cells: Role of the mitochondrial glucocorticoid receptor. Biochimica et Biophysica Acta (BBA) - Molecular Cell Research 1813, 1814–1821 (2011).

12. R. Motavalli et al., The clinical significance of the glucocorticoid receptors: Genetics and epigenetics. J Steroid Biochem Mol Biol 213, 105952 (2021).

13. H. E. Lapp, A. A. Bartlett, R. G. Hunter, Stress and glucocorticoid receptor regulation of mitochondrial gene expression. J Mol Endocrinol 62, R121–r128 (2019).

14. S. M. Smith, W. W. Vale, The role of the hypothalamic-pituitary-adrenal axis in neuroendocrine responses to stress. Dialogues Clin Neurosci 8, 383–395 (2006).

15. G. E. Miller, S. Cohen, A. K. Ritchey, Chronic psychological stress and the regulation of pro-inflammatory cytokines: a glucocorticoid-resistance model. Health Psychol 21, 531–541 (2002).

16. M. Ciccarelli, D. Sorriento, E. Coscioni, G. Iaccarino, G. Santulli, “Chapter 11 - Adrenergic Receptors” in Endocrinology of the Heart in Health and Disease, J. C. Schisler, C. H. Lang, M. S. Willis, Eds. (Academic Press, 2017), https://doi.org/10.1016/B978-0-12-803111-7.00011-7, pp. 285–315.

17. P. B. Molinoff, α- and β-Adrenergic Receptor Subtypes. Drugs 28, 1–15 (1984).

18. H. Zhang et al., α1- and β3-Adrenergic Receptor–Mediated Mesolimbic Homeostatic Plasticity Confers Resilience to Social Stress in Susceptible Mice. Biological Psychiatry 85, 226–236 (2019).

19. P. H. Thaker et al., Chronic stress promotes tumor growth and angiogenesis in a mouse model of ovarian carcinoma. Nat Med 12, 939–944 (2006).

20. S. Li, R. Weerda, C. Milde, O. T. Wolf, C. M. Thiel, ADRA2B genotype differentially modulates stress-induced neural activity in the amygdala and hippocampus during emotional memory retrieval. Psychopharmacology (Berl) 232, 755–764 (2015).

21. S. Hassan et al., Behavioral stress accelerates prostate cancer development in mice. J Clin Invest 123, 874–886 (2013).

22. J. Stemmelin et al., Stimulation of the β3-Adrenoceptor as a Novel Treatment Strategy for Anxiety and Depressive Disorders. Neuropsychopharmacology 33, 574–587 (2008).

23. J. K. MacCormack et al., β-Adrenergic Contributions to Emotion and Physiology During an Acute Psychosocial Stressor. Psychosom Med 83, 959–968 (2021).

24. M. Picard, B. S. McEwen, Psychological Stress and Mitochondria: A Conceptual Framework. Psychosom Med 80, 126–140 (2018).

25. M. Picard, B. S. McEwen, E. S. Epel, C. Sandi, An energetic view of stress: Focus on mitochondria. Frontiers in Neuroendocrinology 49, 72–85 (2018).

26. S. M. Graves et al., Dopamine metabolism by a monoamine oxidase mitochondrial shuttle activates the electron transport chain. Nature Neuroscience 23, 15–20 (2020).

27. M. Karlsson et al., A single-cell type transcriptomics map of human tissues. Sci Adv 7 (2021).

28. E. Sjöstedt et al., An atlas of the protein-coding genes in the human, pig, and mouse brain. Science 367 (2020).

29. G. Consortium, The GTEx Consortium atlas of genetic regulatory effects across human tissues. Science 369, 1318–1330 (2020).

30. M.-S. Kim et al., A draft map of the human proteome. Nature 509, 575–581 (2014).

31. S. C. Bodine, J. D. Furlow, Glucocorticoids and Skeletal Muscle. Adv Exp Med Biol 872, 145–176 (2015).

32. T. Braun, J. R. Challis, J. P. Newnham, D. M. Sloboda, Early-Life Glucocorticoid Exposure: The Hypothalamic-Pituitary-Adrenal Axis, Placental Function, and Long-term Disease Risk. Endocrine Reviews 34, 885–916 (2013).

33. J. Exton, C. Park, Control of gluconeogenesis in liver: II. Effects of glucagon, catecholamines, and adenosine 3′, 5′-monophosphate on gluconeogenesis in the perfused rat liver. Journal of Biological Chemistry 243, 4189–4196 (1968).

34. S. Rath et al., MitoCarta3.0: an updated mitochondrial proteome now with sub-organelle localization and pathway annotations. Nucleic Acids Res 49, D1541–d1547 (2021).

35. G. Sturm et al., A multi-omics longitudinal aging dataset in primary human fibroblasts with mitochondrial perturbations. Scientific Data 9, 751 (2022).

36. G. Sturm et al., Human aging DNA methylation signatures are conserved but accelerated in cultured fibroblasts. Epigenetics 14, 961–976 (2019).

37. K. L. McLaughlin et al., Novel approach to quantify mitochondrial content and intrinsic bioenergetic efficiency across organs. Scientific Reports 10, 17599 (2020).

38. M. Picard, O. S. Shirihai, Mitochondrial signal transduction. Cell Metabolism 34, 1620–1653 (2022).

39. M. Picard, C. Trumpff, Y. Burelle, Mitochondrial Psychobiology: Foundations and Applications. Curr Opin Behav Sci 28, 142–151 (2019).

40. N. Bobba-Alves, R.-P. Juster, M. Picard, The Energetic Cost of Allostasis and Allostatic Load. Psychoneuroendocrinology https://doi.org/10.1016/j.psyneuen.2022.105951, 105951 (2022).

41. C. Trumpff et al., Stress and circulating cell-free mitochondrial DNA: A systematic review of human studies, physiological considerations, and technical recommendations. Mitochondrion 59, 225–245 (2021).

42. F. S. Dhabhar, Effects of stress on immune function: the good, the bad, and the beautiful. Immunol Res 58, 193–210 (2014).

43. R. McCarty, “Chapter 4 - The Fight-or-Flight Response: A Cornerstone of Stress Research” in Stress: Concepts, Cognition, Emotion, and Behavior, G. Fink, Ed. (Academic Press, San Diego, 2016), https://doi.org/10.1016/B978-0-12-800951-2.00004-2, pp. 33–37.

44. R. Glaser, J. K. Kiecolt-Glaser, Stress-induced immune dysfunction: implications for health. Nature Reviews Immunology 5, 243–251 (2005).

45. J. P. Godbout, R. Glaser, Stress-Induced Immune Dysregulation: Implications for Wound Healing, Infectious Disease and Cancer. Journal of Neuroimmune Pharmacology 1, 421–427 (2006).

46. D. A. Padgett, R. Glaser, How stress influences the immune response. Trends in Immunology 24, 444–448 (2003).

47. E. P. Davis, C. A. Sandman, The Timing of Prenatal Exposure to Maternal Cortisol and Psychosocial Stress Is Associated With Human Infant Cognitive Development. Child Development 81, 131–148 (2010).

48. L. A. Grisanti et al., α-Adrenergic Receptors Positively Regulate Toll-Like Receptor Cytokine Production from Human Monocytes and Macrophages. Journal of Pharmacology and Experimental Therapeutics 338, 648–657 (2011).

49. D. Beis et al., The Role of Norepinephrine and α-Adrenergic Receptors in Acute Stress-Induced Changes in Granulocytes and Monocytes. Psychosomatic Medicine 80, 649–658 (2018).

50. N. Bobba-Alves et al., Chronic Glucocorticoid Stress Reveals Increased Energy Expenditure and Accelerated Aging as Cellular Features of Allostatic Load. bioRxiv https://doi.org/10.1101/2022.02.22.481548 (2022).

51. E. Morava, T. Kozicz, Mitochondria and the economy of stress (mal)adaptation. Neurosci Biobehav Rev 37, 668–680 (2013).

52. A. M. Gumpp et al., Childhood maltreatment is associated with changes in mitochondrial bioenergetics in maternal, but not in neonatal immune cells. Proc Natl Acad Sci U S A 117, 24778–24784 (2020).

53. W. Timp, G. Timp, Beyond mass spectrometry, the next step in proteomics. Sci Adv 6, eaax8978 (2020).

54. L. Brydon, K. Magid, A. Steptoe, Platelets, coronary heart disease, and stress. Brain, Behavior, and Immunity 20, 113–119 (2006).

55. M. Uhlén et al., Proteomics. Tissue-based map of the human proteome. Science 347, 1260419 (2015).

56. Anonymous (2021) GTEx Portal. in Dataset Summary of Analysis Samples: V8 Analysis Summary (Broad Institute of MIT and Harvard).

57. Anonymous (The Human Protein Atlas. in Assays & Annotation.

58. J.-C. Goulet-Pelletier, D. Cousineau, A review of effect sizes and their confidence intervals, Part I: The Cohen’s d family. 14, 242–265 (2018).

